# Antibiotic resistance detection is essential for gonorrhoea point-of-care testing: A mathematical modelling study

**DOI:** 10.1101/123620

**Authors:** Stephanie M. Fingerhuth, Nicola Low, Sebastian Bonhoeffer, Christian L. Althaus

## Abstract

Antibiotic resistance is threatening to make gonorrhoea untreatable. Point-of-care (POC) tests that detect resistance promise individually tailored treatment, but might lead to more treatment and higher levels of resistance. We investigate the impact of POC tests on antibiotic-resistant gonorrhoea. We used data about the prevalence and incidence of gonorrhoea in men who have sex with men (MSM) and heterosexual men and women (HMW) to calibrate a mathematical gonorrhoea transmission model. With this model, we simulated four clinical pathways for the diagnosis and treatment of gonorrhoea: POC test with (POC + R) and without (POC − R) resistance detection, culture, and nucleic acid amplification tests (NAATs). We calculated the proportion of resistant infections, cases averted after 5 years, and compared how fast resistant infections spread in the populations. The proportion of resistant infections after 30 years is lowest for POC + R (median MSM: 0.18%, HMW: 0.12%), and increases for culture (MSM: 1.19%, HWM: 0.13%), NAAT (MSM: 100%, HMW: 99.27%), and POC − R (MSM: 100%, HMW: 99.73%). NAAT leads to 36 366 (median MSM) and 1 228 (median HMW) observed cases after 5 years. When compared with NAAT, POC + R results in most cases averted after 5 years (median MSM: 3 353, HMW: 118 per 100 000 persons). POC tests that detect resistance with intermediate sensitivity slow down resistance spread more than NAAT. POC tests with very high sensitivity for the detection of resistance are needed to slow down resistance spread more than using culture. POC with high sensitivity to detect antibiotic resistance can keep gonorrhoea treatable longer than culture or NAAT. POC tests without reliable resistance detection should not be introduced because they can accelerate the spread of antibiotic-resistant gonorrhoea.

## Introduction

Antibiotic resistance is a major challenge for the management of gonorrhoea globally: extended-spectrum cephalosporins are the last antibiotic class remaining for empirical treatment of gonorrhoea [1, 2], and 42 countries have already reported *Neisseria gonorrhoeae* strains with decreased susceptibility against them [2]. The first strain with high-level resistance to recommended combination therapy with ceftriaxone and azithromycin was recently described [3]. With an estimated 78 million new gonorrhoea cases each year [4], new control strategies are urgently needed before gonorrhoea becomes untreatable.

Conventional diagnostic tests for gonorrhoea, nucleic acid amplification tests (NAATs) and culture, are not sufficient to control antibiotic resistance. Commercially available NAATs, the most commonly used diagnostic gonorrhoea tests in high income countries, cannot detect antibiotic resistance [5, 6]. Culture of *N. gonorrhoeae* can be used to determine antibiotic resistance profiles, but reliable results depend on stringent collection and transport of specimens [7]. Both tests need several days to deliver results in routine use. While symptomatic gonorrhoea patients usually receive empirical treatment at their first visit, asymptomatic patients might have to return for treatment. Loss to follow up and further spread of resistant infections can result.

Point-of-care (POC) tests promise to help control antibiotic resistance [8]. POC tests provide results rapidly and allow informed clinical decisions about treatment at the first visit of a patient. POC tests therefore reduce the time to treatment and avoid loss to follow up. A modelling study suggested that POC tests can reduce gonorrhoea prevalence if no antibiotic resistance is present in the population [9]. Though not yet commercially available [8], POC tests that detect resistance promise to reduce the use of antibiotics [10] and to spare last-line antibiotics through individually tailored treatment [11, 12]. One modelling study illustrated that individualised treatment could slow down the spread of resistance as much as combination therapy [13]. However, reduced time to treatment and increased follow up with POC tests might increase the rate of gonorrhoea treatment. Since higher treatment rates can lead to faster spread of resistance [14, 15], POC tests might increase resistance levels. We extended a previously developed mathematical model of gonorrhoea transmission [15] to compare the effects of current conventional tests, culture and NAAT, with POC tests that reduce time to treatment and loss to follow up. We investigated the potential impact of POC tests on resistance and on the number of gonorrhoea cases for a population at high risk of infection [16], men who have sex with men (MSM), and a population at lower risk of infection, heterosexual men and women (HMW).

## Methods

We developed a mathematical model that describes transmission of antibiotic-sensitive and -resistant gonorrhoea, clinical pathways for diagnostic testing with culture, NAAT or POC, and treatment with first- and second-line antibiotics (Supplementary Material: Section Model). Here we describe the model focusing on testing and treatment of gonorrhoea (Fig. 1, Table 1).

**Figure 1.**
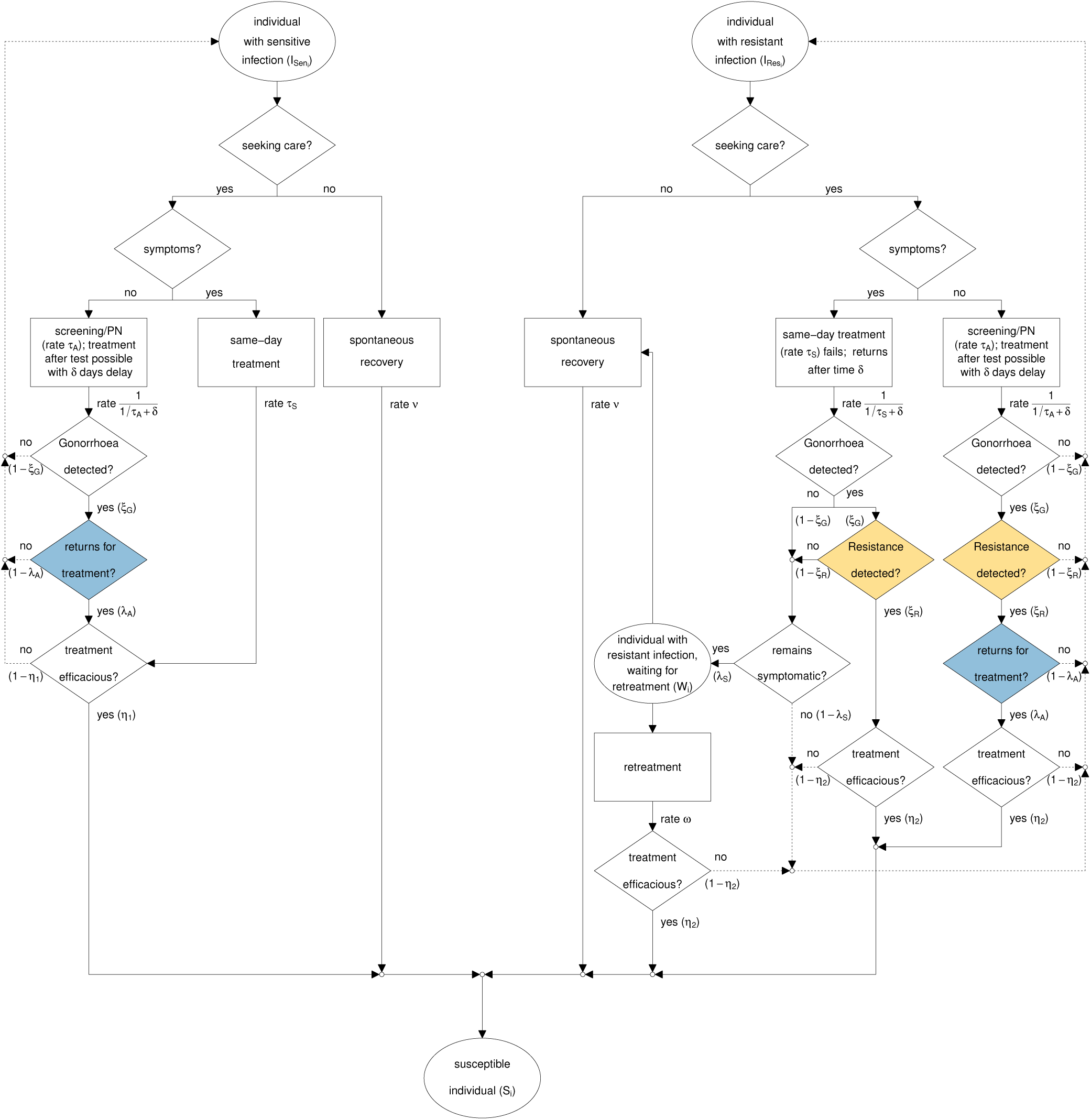
Testing and treatment of gonorrhoea infections. Dashed arrows indicate that individuals remain infected. In the nucleic acid amplification (NAAT) and point-of-care without resistance detection (POC − R) scenario, “Resistance detected?” (yellow) defaults to “no”. In all point-of-care scenarios, “returns for treatment?” (blue) defaults to “yes”. In the culture scenario, the flowchart is followed as shown. PN: partner notification.

**Figure 2.**
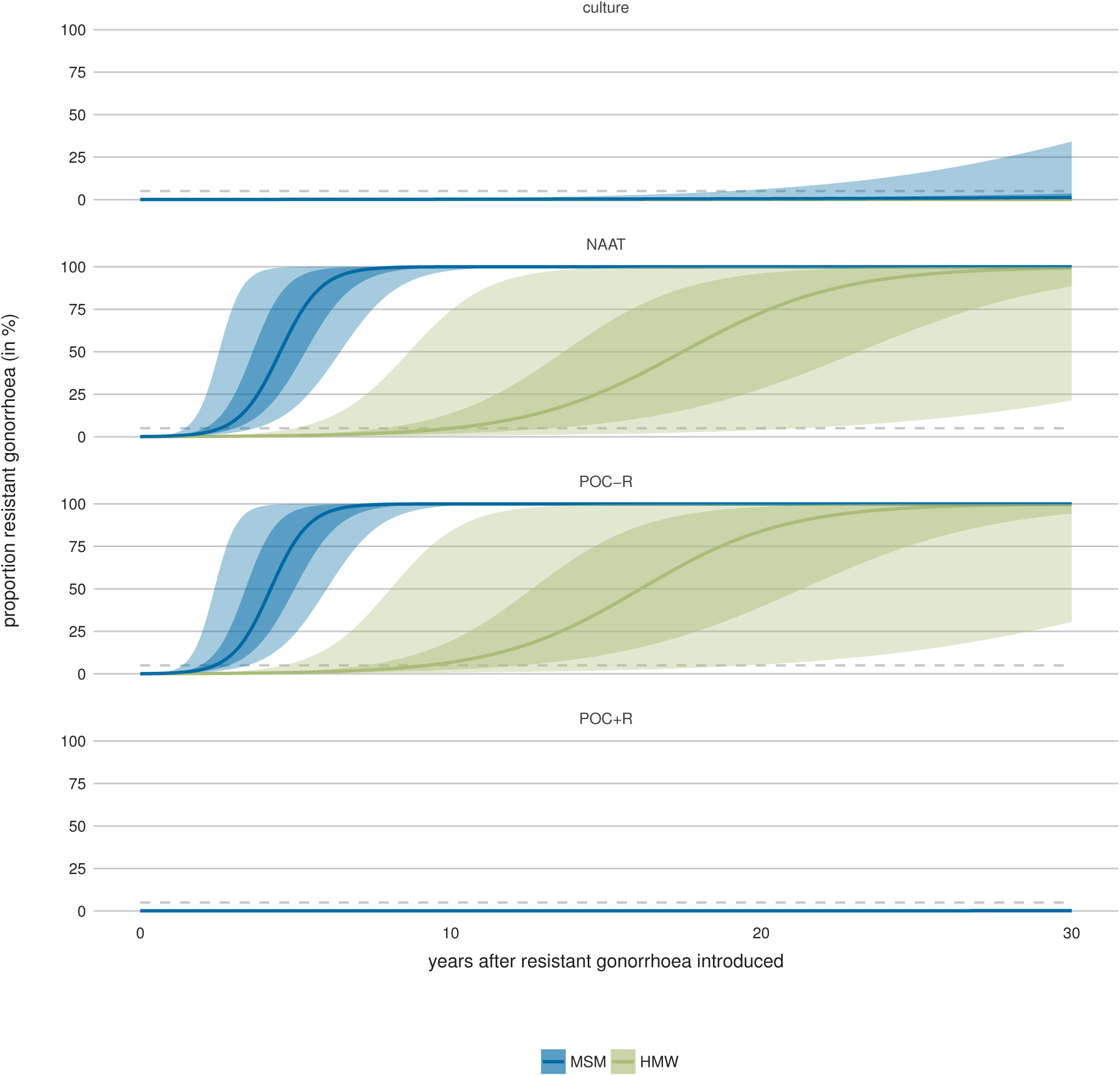
Proportion of resistant gonorrhoea infections for each testing scenario. The continuous lines give the median proportion of resistant infections over all simulations. Shaded areas indicate that 50% or 95% of all simulations lie within this range. MSM: men who have sex with men, HMW: heterosexual men and women. The proportion of resistant infections remains lowest when point-of-care with resistance detection (POC + R) is used, followed by culture. The proportion of resistant infections exceeds the 5% threshold (dashed lines) marginally earlier with point-of-care without resistance detection (POC − R) than with the nucleic acid amplification test (NAAT).

**Table 1.**
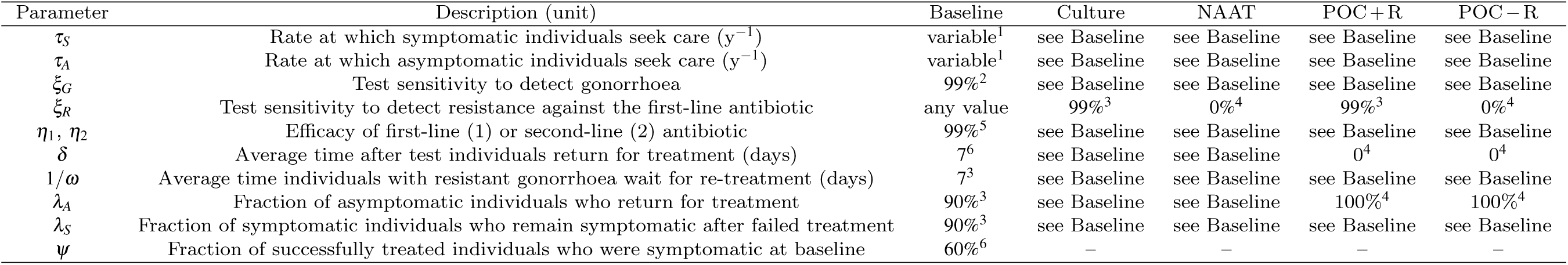
Gonorrhoea testing and treatment parameters and their default values. Unless a value is set by definition, all values listed are default values and are varied in sensitivity analyses. Baseline: resistance-free scenario (corresponds to scenario where culture or nucleic acid amplification test (NAAT) is used; *ξ_R_* can take any value since there is no resistance to detect). Culture, NAAT, Point-of-care (POC) with resistance detection (POC + R) and without resistance detection (POC − R) refer to scenarios after resistance is introduced. Sources for parameters: ^1^Derived, ^2^[17], ^3^Assumption, ^4^by definition, ^5^[18, 19], ^6^[20].

### Basic model structure

The model is based on our previously published compartmental model of gonorrhoea transmission and resistance spread [15]. The model describes a population with two sexual activity classes *i* ∈*C*, where *C* = {*L*,*H*} indicates that there are two sexual activity classes *L* and *H* with low and high partner change rates. The model incorporates sexual mixing between the sexual activity classes, sexual behaviour change, migration in and out of the population, and gonorrhoea transmission. Individuals in the population can be susceptible to infection, *S_i_*, infected with antibiotic-sensitive gonorrhoea, *I_Sen_i__*, infected with gonorrhoea resistant to the first-line antibiotic, *I_Res_i__*, or infected with gonorrhoea resistant to the first-line antibiotic and waiting for re-treatment, *W_i_*. Depending on the parameters for sexual behaviour, transmission, and gonorrhoea natural history (Supplementary Material: Table S2), the model describes a population of men who have sex with men (MSM) or heterosexual men and women (HMW).

### Gonorrhoea testing and treatment

#### Antibiotic-sensitive gonorrhoea

Individuals infected with antibiotic-sensitive gonorrhoea, *I_Sen_i__*, (Fig. 1, left) can recover spontaneously at rate *v* or seek care. Symptomatic care-seekers receive treatment on the same day at rate *τ_s_*. Asymptomatic care-seekers, i.e. those who are screened for gonorrhoea or were notified through an infected partner, are tested at rate *τ_A_*. Gonorrhoea is detected with sensitivity *ξ_G_*. On average, a fraction *λ_A_* of asymptomatic individuals returns for treatment after *δ* days. The treatment rate for asymptomatic individuals is approximated by 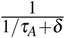, the inverse of the average time until individuals are tested, 1/*τ_A_*, and the time until they return for treatment, *δ*. Both symptomatic and asymptomatic individuals are treated with a first-line antibiotic that has treatment efficacy *η*_1_. We assumed that individuals whose treatment was inefficacious remain infected and do not seek care again immediately. This assumption reflects the notion that treatment failure of antibiotic-sensitive gonorrhoea is most likely to occur in pharyngeal infections, which are usually asymptomatic [21].

#### Antibiotic-resistant gonorrhoea

Individuals infected with gonorrhoea resistant to the first-line antibiotic, *I_Res_i__*, (Fig. 1, right) can also recover spontaneously at rate *v*. Asymptomatic care-seekers that return for treatment (fraction *λ_A_*) receive treatment with the second-line antibiotic at rate 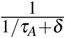 if both gonorrhoea (sensitivity *ξ_G_*) and resistance (sensitivity *ξ_R_*) are detected. Symptomatic care-seekers receive the first-line antibiotic as treatment on the same day, but remain infected due to resistance and return for treatment after *δ* days. At their second visit, symptomatic care-seekers receive the second-line antibiotic if both gonorrhoea (sensitivity *ξ_G_*) and resistance (sensitivity ξ*_R_*) are detected. If either test fails, they do not receive the second-line antibiotic. If they remain symptomatic (fraction *λ_s_*), they wait for re-treatment in compartment *W_i_*, where they either receive re-treatment with the second-line antibiotic at rate *ω* or recover spontaneously at rate *ν*. The assumption that re-treatment occurs with the second-line antibiotic follows recommendations from the World Health Organization (WHO) [16] and the Centers for Disease Control (CDC) [22] to obtain a specimen for culture-based antibiotic resistance testing at a patient’s second visit. The second-line antibiotic has efficacy *η*_2_; individuals whose treatment is inefficacious remain infected and can recover spontaneously or seek care at a later point. De novo resistance to the first-line antibiotic or resistance to the second-line antibiotic are not considered in the model.

### Testing scenarios

The model allowed us to simulate clinical pathways for gonorrhoea detection with culture, NAAT, and POC tests by adapting the parameters *δ, λ_A_*, and ξ*_R_* (Table 2). For culture, test results are not available immediately (*δ*_culture_ > 0), resistance can be detected (*ξ*_*R*, culture_ > 0), and asymptomatic infected individuals might not return for treatment (*λ*_*A*, culture_ < 1). For NAAT, test results are not available immediately (*δ*_NAAT_ > 0), resistance cannot be detected (*ξ*_*R*, NAAT_ = 0), and asymptomatic infected individuals might not return for treatment (*λ*_*A*, NAAT_ < 1). For POC, test results are available immediately (*δ*_POC_ = 0), all individuals are followed up (*λ*_*A*, POC_ = 1), and thus all individuals are treated at the first visit. We explore the impact of a POC test with (*ξ*_*R*, POC_ > 0, POC + R) and without resistance detection (*ξ*_*R*, POC_ = 0, POC − R); we use the term “POC” alone when *ξ*_*R*, POC_ is variable.

**Table 2.** Conditions on parameters for different testing scenarios. NAAT: nucleic acid amplification test, POC: point-of-care test (with or without resistance detection), POC + R: POC test with resistance detection, POC − R: POC test without resistance detection.

| | $\delta$ | $\lambda_A$ | $\xi_R$ |
| --- | --- | --- | --- |
| Culture | $> 0$ | $< 1$ | $> 0$ |
| NAAT | $> 0$ | $< 1$ | $= 0$ |
| POC | $= 0$ | $= 1$ | $\geq 0$ |
| POC + R | $= 0$ | $= 1$ | $> 0$ |
| POC – R | $= 0$ | $= 1$ | $= 0$ |

### Impact measures

We evaluated the impact of a testing scenario by calculating the proportion of resistant infections among all infections, observed cases averted, and the rate at which resistance spreads, compared with another testing scenario. We measured the proportion of resistant infections up to 30 years after introduction of resistance into the resistance-free baseline scenario. If applicable, we also calculated the time until resistance levels reached 5%, the level above which an antibiotic should not be used for empirical gonorrhoea treatment [18]. We defined observed cases averted as the difference between the cumulative incidence of observed (i.e. diagnosed and successfully treated at baseline; fraction *ϕ* [15]) cases using NAAT and the cumulative incidence of observed cases using culture or POC tests. We calculated the observed cases averted 5 years after the introduction of resistance. The rate at which resistance spreads describes how fast resistant infections replace sensitive infections in a human population [15]. We calculated the ratio of the rate of resistance spread between POC with different test sensitivities to detect resistance (*ξ*_*R*, POC_) and culture or NAAT scenarios (Supplementary Material: Section Rate of resistance spread and ratio of resistance spread). If the ratio of the rate of resistance spread is > 1, resistance spreads faster when using POC tests compared with other tests. If the ratio is < 1, resistance spreads slower when using POC tests compared with other tests.

### Parameters

We used the parameters describing sexual behaviour, gonorrhoea transmission, natural history, and treatment from our previous model [15]. There, we estimated sexual behaviour parameters from the second British National Survey of Sexual Attitudes and Lifestyles (Natsal-2), which is a nationally representative population-based survey [23]. We calibrated all other parameters to yield prevalence and incidence rates within empirically observed ranges (Table 3 and 4). For this study, we used a subset of 1 000 calibrated parameter sets from the previous study. For each calibrated parameter set, we derived the care seeking rate of asymptomatic (*τ_A_*) and symptomatic (*τ_s_*) individuals using the fraction of successfully treated individuals who were symptomatic at baseline *ϕ* (Supplementary Material: Section Derivation of *τ_A_* and *τ_S_*). We set default values for the testing and treatment parameters *ψ, ξ_G_, ξ_R_, η*_1_, *η*_2_, *δ, ω, λ_A_* and *λ_S_* guided by literature (Table 1).

**Table 3.** Gonorrhoea prevalence and incidence in baseline scenario (before resistance introduced) for MSM. The prevalence and incidence ranges used for calibration for men who have sex with men (MSM) were based on the Health in Men Study in Australia [24]. The baseline median and interquartile range (IQR) are based on the simulation results of 1000 calibrated parameter sets. The upper and lower bound of the calibration range for the low and high sexual activity class were set to the lower and upper bound for the total population. The calibration is detailed in [15].

|  | Range used for calibration | Baseline median (IQR) |
| --- | --- | --- |
| Prevalence low activity class (%) | 0 – 2.79 | 0.59 (0.42 – 0.79) |
| Prevalence high activity class (%) | 1.19 – 100 | 27.64 (23.25 – 31.91) |
| Prevalence total population (%) | 1.19 – 2.79 | 2.09 (1.69 – 2.43) |
| Incidence total population (100000 persons<sup>-1</sup> y<sup>-1</sup>) | 5880 – 7190 | 6493.49 (6192.89 – 6842.70) |

**Table 4.** Gonorrhoea prevalence and incidence in baseline scenario (before resistance introduced) for HMW. The prevalence and incidence ranges used for calibration for Heterosexual men and women (HMW) were based on the National Health and Nutrition Examination Survey [25] and surveillance data [26], both from CDC. The baseline median and interquartile range (IQR) are based on the simulation results of 1000 calibrated parameter sets. The upper and lower bound of the calibration range for the low and high sexual activity class were set to the lower and upper bound for the total population. The calibration is detailed in [15].

|  | Range used for calibration | Baseline median (IQR) |
| --- | --- | --- |
| Prevalence low activity class (%) | 0 – 0.38 | 0.12 (0.09 – 0.15) |
| Prevalence high activity class (%) | 0.16 – 100 | 2.14 (1.71 – 2.60) |
| Prevalence total population (%) | 0.16 – 0.38 | 0.25 (0.21 – 0.3) |
| Incidence total population (100000 persons<sup>-1</sup> y<sup>-1</sup>) | 120 – 360 | 222.13 (172.19 – 283.54) |

### Sensitivity Analyses

We performed sensitivity analysis to confirm that our model results are robust in scenarios with different properties of tests (*ξ_G_, ξ_R_*), antibiotics (*η*_1_, *η*_2_), and populations and clinics (*δ, ω, λ_A_, λ_S_*). First, we performed sensitivity analyses of the number of observed cases averted with regard to changes in both the fraction of asymptomatic individuals who return for treatment at baseline (*λ_A_*) and fraction of successfully treated individuals who were symptomatic at baseline (*ψ*) (Fig. 3), as well as to changes in single testing and treatment parameters (*ξ_G_, ξ_R_, λ_A_, λ_S_, ψ, δ, ω*, Supplementary Material: Figures S3-S9). Second, we evaluated the sensitivity of the ratio of resistance spread with regard to changes in the test sensitivity to detect resistance against the first-line antibiotic when using POC (*ξ*_*R*, POC_), the fraction of asymptomatic individuals who return for treatment at baseline (*λ*_*A*, baseline_) and the fraction of successfully treated individuals who were symptomatic at baseline (*ψ*, Fig. 4 and 5). Third, we tested the sensitivity of our model results to the assumption that the test sensitivity to detect *N. gonorrhoeae* (*ξ_G_*) is 99% for culture testing. For this, we simulated an alternative baseline scenario where only culture, with a test sensitivity to detect *N. gonorrhoeae* (*ξ_G_*) of 90%, is used (all other parameters as in Table 1, Supplementary Material: Figures S10-S12).

**Figure 3.**
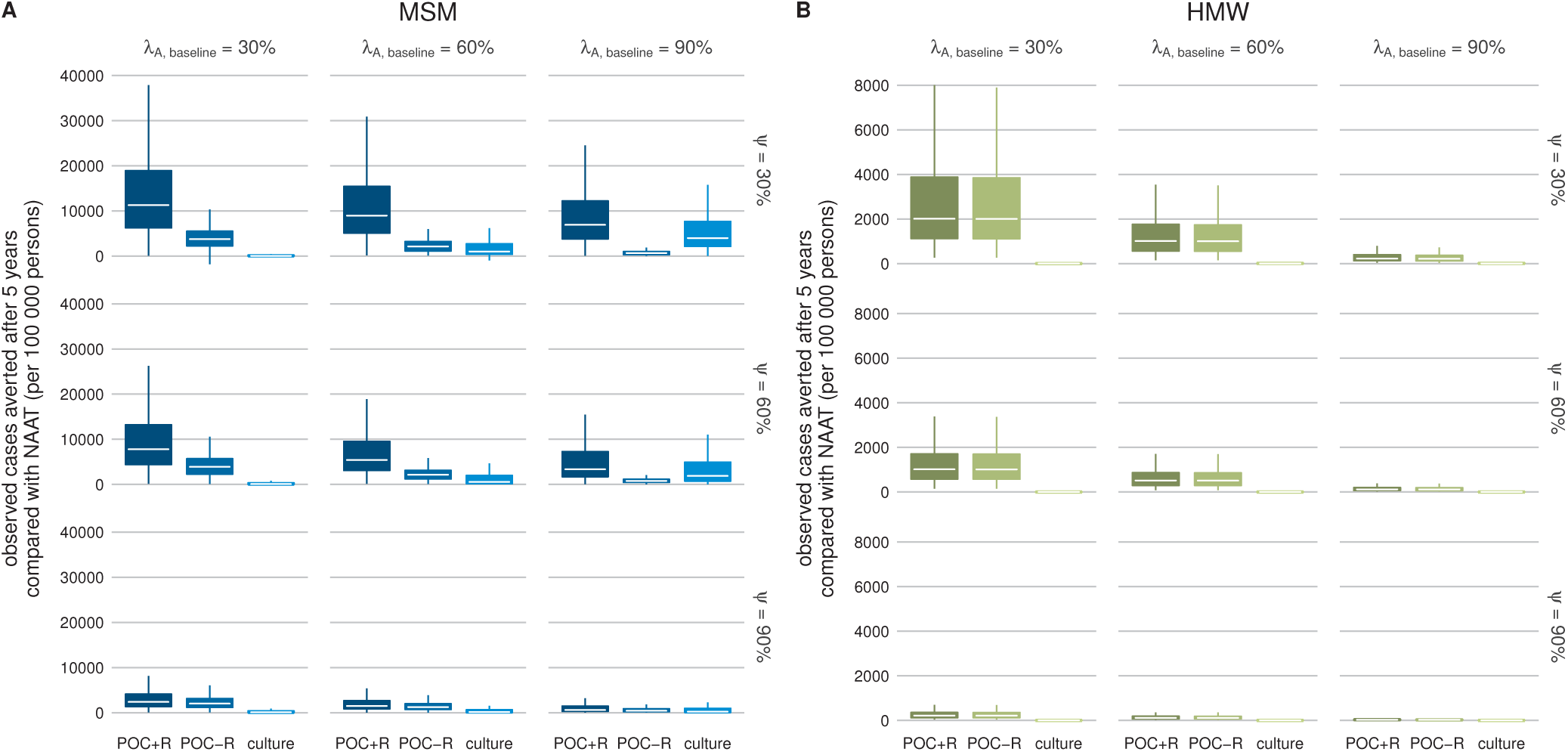
Two-dimensional sensitivity analysis of observed cases averted (per 100000 persons) after 5 years. The sensitivity analysis is performed with respect to the fraction of asymptomatic individuals who return for treatment at baseline (*λ*_*A*, baseline_) and the fraction of successfully treated individuals who were symptomatic at baseline (*ψ*), for (A) men who have sex with men (MSM) and (B) heterosexual men and women (HMW). The central right plot of each panel shows the default scenario (*λ*_*A*, baseline_ = 90%, *ψ* = 60%). NAAT: nucleic acid amplification test, POC + R: point-of-care test (POC) with resistance detection, POC − R: POC without resistance detection. Lower/upper bound of the box indicate first/third quartiles, bar in box indicates median, whiskers span 1.5 times interquartile range. Outliers not shown for more clarity.

### Simulation

For each parameter set, we first simulated a resistance-free baseline scenario where either culture or NAAT is used (*δ* > 0, *λ_A_* < 1). We simulated the baseline scenario until it reached equilibrium using the function *runsteady* from the R language and software environment for statistical computing [27] package rootSolve [28]. Next, we introduced resistant strains by converting 0.1% of all sensitive infections into resistant infections. We then set the parameter *ξ_R_* to reflect the different testing scenarios (culture, NAAT, POC + R or POC − R). For POC tests, we additionally set *δ* = 0 and *λ_A_* = 1. Finally, we simulated the model using the function *lsoda* from the R package deSolve [29].

## Results

### Proportion of resistant infections

We determined the proportion of gonorrhoea infections resistant to the first-line antibiotic for up to 30 years after the introduction of resistance (Fig. 2). The proportion of resistant infections remains lowest when POC + R is used (MSM: median 0.18% after 30 years, interquartile range (IQR) 0.17 − 0.21%; HMW: 0.12%, 0.11 − 0.12%). The proportion of resistant infections also remains low with culture (MSM: 1.19%, 0.68 − 3.59%, HWM: 0.13%, 0.12 − 0.15%). In contrast, resistant infections largely replace sensitive infections after 30 years using NAAT (MSM: 100%, 100 −100%, HMW: 99.27%, 88.54 − 99.97%) and POC − R (MSM: 100%, 100 − 100%, HMW: 99.73%, 94.30 − 99.99%). The proportion of resistant infections exceeds the 5% resistance threshold (Fig. 2, dashed line) marginally earlier when POC − R is used (MSM: median < 2.42, IQR 2.00 − 2.92 years, HMW: < 9.25, 7.25 − 12.25 years) than when NAAT is used (MSM: < 2.58, 2.08 − 3.08 years, HMW: < 10.08, 7.83 − 13.33 years). Overall, POC + R performs best in keeping the proportion resistant infections low and POC − R performs worst.

### Observed cases averted

We calculated the observed cases averted (per 100 000 persons) after 5 years using culture, POC + R or POC − R in comparison with NAAT (Fig. 3). For the default values (*λ*_*A*, baseline_ = 90%, *ψ* = 60%), using NAAT leads to a median of 36366 (IQR 33789 − 39692) observed cases after 5 years for MSM and 1228 (927 − 1610) for HMW. Culture averts 1876 (740 − 4919) cases in MSM and 3 (1 − 7) in HMW compared with NAAT. POC + R averts even more cases than culture in both MSM (3353, 1697 − 7259) and HMW (118, 69 − 198). POC − R averts less cases than culture in MSM (772, 452 − 1119), but about the same as POC + R in HMW (115, 68 − 190).

For culture, increasing the fraction of asymptomatic individuals who return for treatment at baseline (*λ*_*A*, baseline_) and decreasing the fraction of successfully treated individuals who were symptomatic at baseline (*ψ*) increases the median number of observed cases averted. For POC + R, decreasing *λ*_*A*, baseline_ and decreasing *ψ* leads to an increase in the median observed cases averted. For POC − R, decreasing *λ*_*A*, baseline_ and the intermediate value of *ψ* results in an increase in median averted cases. For all combinations of *λ*_*A*, baseline_ and *ψ* in both MSM and HMW, POC + R averts more cases at the median than culture. This result is robust to changes in single testing and treatment parameters (Supplementary Material: Figures S3-S9).

### Ratio of resistance spread

We determined the ratio of the rate of resistance spread between POC and culture (Fig. 4) and POC and NAAT (Fig. 5). For the default values (*ξ*_*R*, culture_ = 99%, *ξ*_*R*, NAAT_ = 0%, *ξ*_*R*, POC_ = 99%, *λ*_*A*, baseline_ = 90%, *ψ* = 60%), resistance spreads more slowly with POC compared with culture or NAAT. Decreasing the test sensitivity to detect resistance of POC (*ξ*_*R*, POC_) can result in a faster spread of resistance with POC. A slight decrease in *ξ*_*R*, POC_ to 80-95% already leads to faster resistance spread with POC compared with culture. In contrast, only very low values of *ξ*_*R*, POC_ result in a faster resistance spread for POC compared with NAAT.

**Figure 4.**
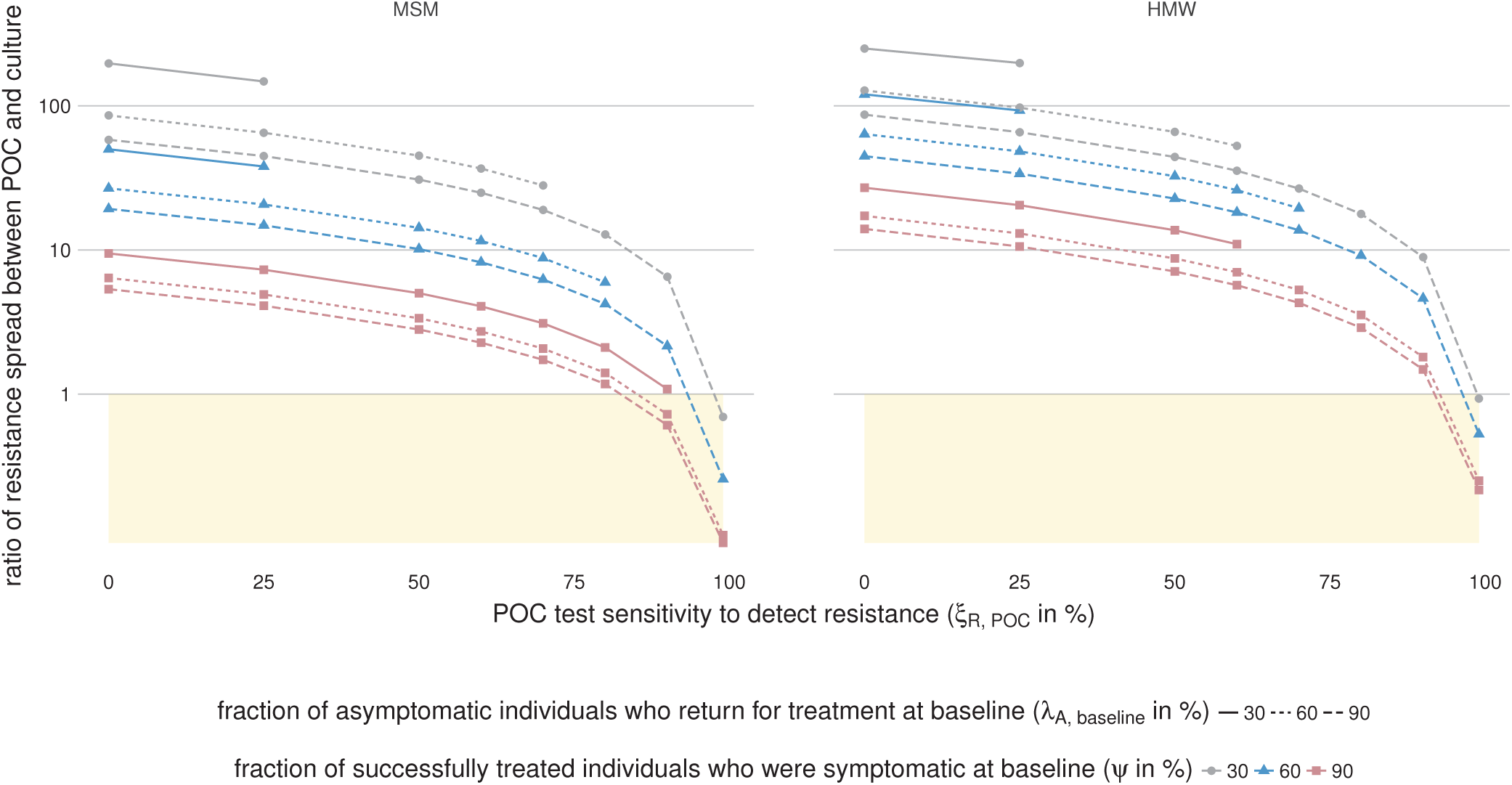
Ratio of resistance spread between point-of-care test (POC) and culture. Shown are the ratios of resistance spread for men who have sex with men (MSM) and heterosexual men and women (HMW) for *ξ*_*R*, culture_ = 99% and different values of *ξ*_*R*, POC_, *λ*_*A*, baseline_ and *ψ*(POC − R: *ξ*_*R*, poc_ = 0, POC + R: *ξ*_*R*, poc_ > 0). The shaded areas indicate that resistance spread is slower when using POC than when using culture. For the default values (*ξ*_*R*, POC_ = 99%, *λ*_*A*, baseline_ = 90%, *ψ* = 60%), resistance spread is slower when using POC than when using culture. For most other shown values using POC accelerates resistance spread. Each data point gives the median value over 1000 simulations (one per calibrated parameter set). Some calibrated parameter sets lead to extinction of gonorrhoea in the simulation (Supplementary Material: Figure S2). In these simulations, resistance did not spread and the ratio of resistance spread could not be calculated. Data points that would include such simulations were excluded from this figure since they would show the median ratio of resistance spread over less than 1000 simulations. Note that the y-axis is shown in logarithmic scale.

## Discussion

Using a mathematical transmission model, we compared the expected impact of POC tests on gonorrhoea cases and antibiotic resistance with conventional tests, culture and NAAT. We found that POC tests that detect antibiotic resistance avert more gonorrhoea cases than any other test across all simulated settings. Compared with culture and NAAT, POC tests with high sensitivity to detect resistance slow the spread of resistant infections. POC tests with no or low sensitivity to detect resistance accelerate the spread of resistant infections.

We captured the basic principles of the gonorrhoea testing and treatment process for culture, NAAT and POC in a single model structure. The parameters describing the sexual behaviour and the natural history of gonorrhoea were estimated and calibrated in a previous study [15]. The default parameters that describe testing and treatment of gonorrhoea were based on literature values and are measurable. The model results are robust in sensitivity analyses (Fig. 3 and 4, 5, Supplementary Material: 23Figures S3-S12).

**Figure 5.**
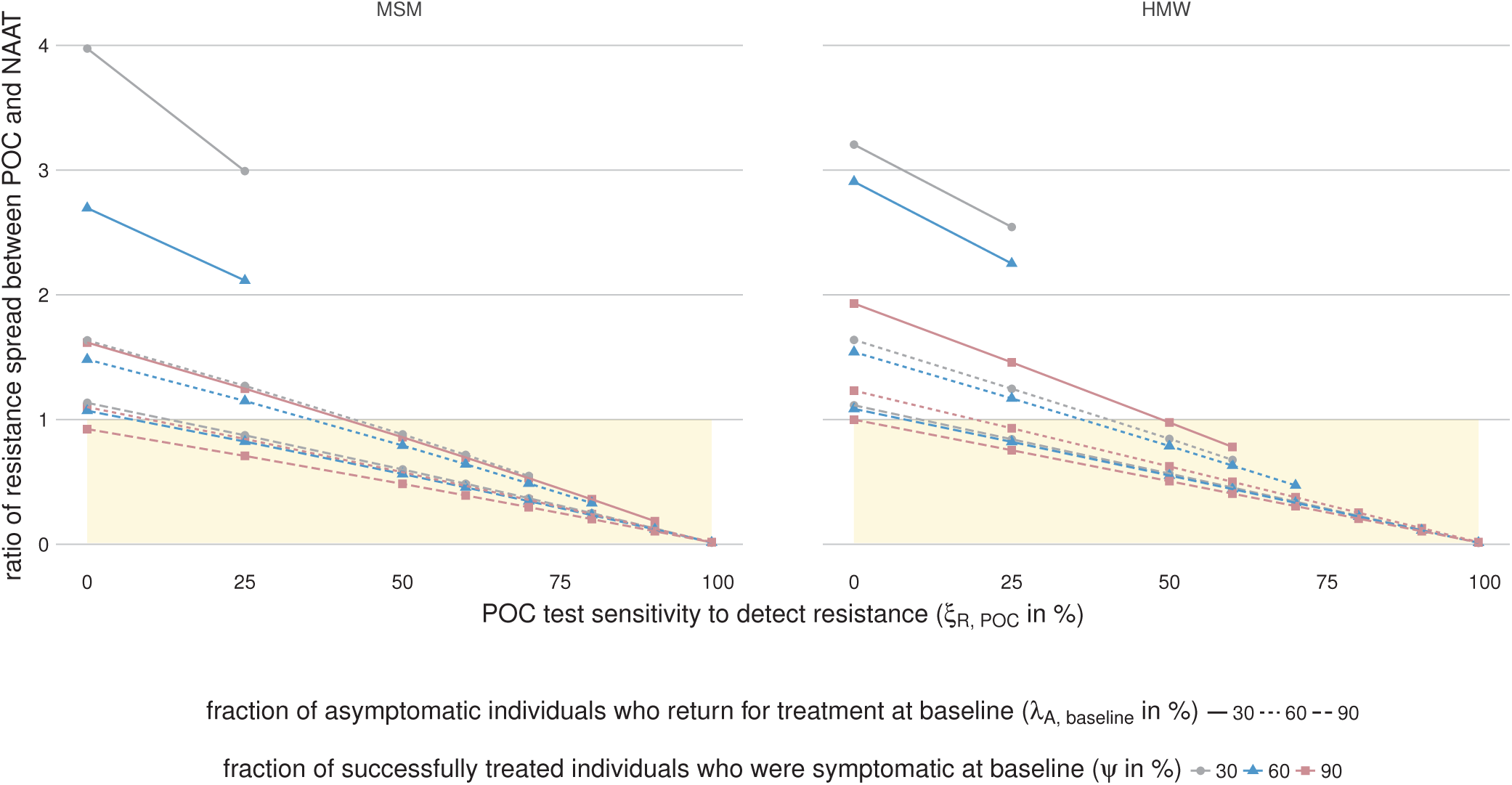
Ratio of resistance spread between point-of-care test (POC) and nucleic acid amplification test (NAAT). Shown are the ratios of resistance spread for men who have sex with men (MSM) and heterosexual men and women (HMW) for *ξ*_*R*, NAAT_ = 0% and different values of *ξ*_*R*, POC_, *λ*_*A*, baseline_ and *ψ* (POC − R: *ξ*_*R*, POC_ = 0, POC + R: *ξ*_*R*, POC_ > 0). The shaded areas indicate that resistance spread is slower when using POC than when using NAAT. For the default values (*ξ*_*R*, POC_ = 99%, *λ*_*A*, baseline_ = 90%, *ψ* = 60%) and most other shown values resistance spread is slower when using POC than when using NAAT. Each data point gives the median value over 1000 simulations (one per calibrated parameter set). Some calibrated parameter sets lead to extinction of gonorrhoea in the simulation (Supplementary Material: Figure S2). In these simulations, resistance did not spread and the ratio of resistance spread could not be calculated. Data points that would include such simulations were excluded from this figure since they would show the median ratio of resistance spread over less than 1000 simulations.

We managed the complexity of our model with the following assumptions: First, we did not consider test specificity. A low test specificity to detect resistance against the first-line antibiotic would result in increased use of the second-line antibiotic, and thus simultaneously decrease the level of resistance against the first-line antibiotic and increase the level of resistance against the second-line antibiotic. Since we focused on resistance against the first-line antibiotic, we could not capture the impact of test specificity appropriately. Second, our model does not include a change in antibiotic recommendations: undetected resistant infections are always treated with the first-line antibiotic, even if all infections in the population are resistant. This clinical pathway increases the average duration of resistant infections and possibly the observed cases. Whilst this is unlikely in high income countries with good antibiotic resistance surveillance, it is not an unrealistic scenario in resource poor settings without surveillance where 71-100% of gonococcal strains are resistant to fluoroquinolones [30]. In our model, MSM have a substantial level of resistant gonorrhoea infections after 5 years using NAAT. We expect that our model overestimates the observed cases using NAAT and the observed cases averted using culture and POC + R compared with a model including antibiotic recommendation change. Third, we considered only treatment with a single antibiotic although current treatment guidelines recommend combination therapy with two antibiotics simultaneously [1, 7]. The model results are fully applicable to treatment with combination therapy if antibiotic-resistant gonorrhoea is interpreted as resistance against both antibiotics used for combination therapy. Fourth, we investigated the effects of one test at a time and did not consider the effects of mixed testing. Our results therefore only show what the ideal effects of each test could be. Fifth, we simplified the testing and treatment process. To better compare the testing scenarios, we did not model care seeking and returning for treatment as separate processes, but approximated the overall treatment rates. In accordance with WHO [16] and CDC recommendations [22], we assumed that re-treatment of resistant infections occurs with the second-line antibiotic because a resistance profile has been determined after the second visit. Finally, for better comparability we assumed that culture, NAAT and POC tests have the same sensitivity to detect gonorrhoea, even though culture has lower sensitivity to detect rectal or pharyngeal gonorrhoea than molecular tests [31]. A lower test sensitivity to detect gonorrhoea, *ξ_G_*, requires a higher care-seeking rate of asymptomatic individuals, *τ_A_*, to obtain the same prevalence and incidence rates. We simulated an alternative scenario (Supplementary Material: Figures S10-S12) where only culture is used at baseline (with *ξ_G_* = 90% for culture and all other values as in Table 1). In this scenario, the proportion of resistant infections after 30 years using culture is higher in MSM (median 3.18%, IQR 1.51 − 11.33%) and the observed cases averted after 5 years using POC + R compared with NAAT is larger (median 4 236, IQR 2161 − 8839 per 100000 persons). Overall the effect of lower test sensitivity to detect gonorrhoea with culture was small.

Currently, there are no commercial POC tests that can detect antibiotic-resistant *N. gonorrhoeae* [8] and there remain challenges for their development. First, molecular POC test that detect resistance need molecular markers that reliably predict phenotypic resistance. So far only markers that predict resistance against some antibiotics are known [8, 32, 33]. Second, diagnostic tests need to deliver results fast to be considered point-of-care. The fastest molecular diagnostic test for gonorrhoea that is commercially available takes 90 minutes [34, 35] which might be too long to wait for some patients. Finally, costs and training requirements for molecular tests have hindered their availability in low income countries so far [36].

This study addresses two key questions for gonorrhoea control and resistance [37]. First, we investigated the potential impact of a POC test that detects antibiotic resistance (POC + R). We found that POC + R can slow resistance spread and reduce the gonorrhoea cases compared with culture or NAAT. The impact of POC + R is particularly strong when the fraction of asymptomatic individuals who return for treatment (*λ*_*A*, baseline_) and the fraction of successfully treated individuals who were symptomatic (*ψ*) were low before POC + R is introduced. However, when the POC test cannot detect resistance (POC − R) the benefits of POC are outweighed by accelerated resistance evolution: because fewer patients are lost to follow up, more patients are treated and more antibiotic treatment selects more strongly for antibiotic resistance. Since resistance cannot be detected, resistance levels increase and fewer cases are averted. Second, we investigated the impact of POC tests in two populations with different levels of risk of gonorrhoea, MSM and HMW. We found that in both populations, POC tests with reliable resistance detection (POC + R) slow down the spread of resistance and avert the most cases. POC tests without resistance detection (POC − R) avert about as many cases as POC + R in HMW, but clearly fewer cases than POC + R in MSM. Since resistance usually spreads faster in MSM [15], the faster spread of resistance caused by POC − R already impacts the cases averted after 5 years in MSM, but not yet in HMW. POC tests that detect resistance reliably are crucial for both populations and both populations need culture-based surveillance of resistance to keep molecular markers for POC resistance detection updated.

This modelling study addresses clinically relevant situations, can be used to help design trials comparing different test strategies and guide the introduction of POC tests in the future. POC tests with high sensitivity to detect resistance can replace culture-based diagnosis in clinical settings, as long as culture-based surveillance of antibiotic resistance is maintained to monitor resistance levels and to determine molecular markers for POC tests. POC tests with lower sensitivities to detect resistance should not replace culture-based diagnosis, but might have some advantages over NAAT. POC test with low or no sensitivity to detect resistance should not be introduced, because POC tests without reliable resistance detection can accelerate the spread of antibiotic-resistant gonorrhoea.

## Author’s contributions

SMF, NL, SB, CLA designed the study. SMF simulated the model. SMF, NL, SB, CLA interpreted the data. SMF wrote the first version of the manuscript. All authors contributed to and approved the final version of manuscript.

